# Assembly-Free Short-Read Metagenomic Maximum Growth Rate Prediction

**DOI:** 10.1101/2025.09.18.677194

**Authors:** Jesse Natarajan, JL Weissman

## Abstract

Microbial maximum growth rates are key parameters in microbially explicit soil biogeochemical models that can also be estimated directly from genomes and metagenomes, therefore enabling model-data synthesis between coarse-grained models and high-resolution sequencing data. A persistent challenge for predicting community-level growth phenotypes in soil metagenomes is that current maximum growth rate prediction models require complete gene sequences from assembled contigs, but the complexity of soil metagenomes often precludes high quality assembly. Here, we develop a set of gene length specific models that are capable of predicting community average maximum growth rates directly from genes inferred from reads, avoiding the need for an assembly step. These models have been implemented in the open source *gRodon* R package for maximum growth rate prediction.

## Introduction

Soil microbial communities play a central role in soil biogeochemical cycling and are key mediators of soil health and ecosystem productivity [1, 2]. As anthropogenic climate change progresses, predicting the functioning of soil microbial communities in changing environments will be increasingly important for understanding climate-soil feedbacks. Yet, the incredible taxonomic and functional diversity of these communities makes them challenging to integrate into earth systems models [3].

One path forward is to measure traits of microbial populations that can then be incorporated as parameters into biogeochemical models (*e.g*., [4, 5]). Yet, the inability to grow most soil microbes in the laboratory raises the need for techniques to analyze the functional capabilities of these organisms that do not require cultivation [6]. The development of ‘omics tools has given us a detailed picture of microbial diversity at a molecular scale, but these complex datasets do not easily translate to global-scale understanding. For example, analysis of gene presence or absence can allow the reconstruction of complex metabolic networks from environmental metagenomes (*e.g*., [7–9]), but the process for converting these descriptions of hypothetical biochemistry to model parameters is not always obvious.

As an alternative, genomic trait prediction approaches permit the inference of complex traits that often have direct model parameter analogs using coding and non-coding features throughout the genome (e.g., [10–12]).

For example, an organism’s maximum growth rate, which is a key parameter in models of microbial community dynamics can be inferred directly from patterns of translation optimization in bacterial, archaeal, and even eukaryotic genomes and metagenomes [10, 13–17]. Highly expressed genes, particularly those encoding ribosomal proteins and other components of the translation machinery, tend to exhibit stronger codon usage bias (CUB), favoring codons that correspond to the most abundant tRNAs in the cell [18–21]. Due to this relationship between codon usage and translation efficiency, CUB is frequently used as a predictor of microbial growth rates [10, 13]. Extending to the community level, it is possible to estimate the average maximum growth rate of an entire microbial community (or prefiltered subset, e.g., by taxonomy, of that community) by applying growth rate prediction tools directly to metagenomes [13, 17]. Together, genomic and metagenomic growth rate predictors present promising opportunities for model-data synthesis between soil metagenomes and earth systems models.

Methodologically, the application of genomic growth potential prediction tools to soil metagenomes is complicated by the fact that short-read metagenomes, still by far the cheapest and most common type of metagenomic sequences for soil microbes, demonstrate sufficiently high sequence diversity to preclude high-quality assembly or binning for most community members [22], especially for low-to-medium coverage metagenomes. While coding sequences can be inferred directly at the read level [23], the application of CUB-based maximum growth rate predictors to coding sequences inferred directly from reads leads to poor quality predictions because CUB estimates are strongly biased when applied to short sequences [17]. This has led to previous claims by one of our authors that assembly is a necessary step for maximum growth rate prediction [17]. Here, we show that this is not the case, and that by building gene length specific models trained on truncated gene sequences we can predict the average maximum growth rate of a microbial community directly from reads without an assembly step at comparable accuracy to predictions made from assembled contigs. We incorporate these length specific models into the open source *gRodon* R package for maximum growth rate prediction.

## Results & Discussion

### Measures of Codon Usage Bias are Biased for Very Short Sequences

We assessed how sequence length influenced CUB estimates for 8,300 genome assemblies from bacteria and archaea in the original *gRodon* training set [10, 24]. Specifically, for each genome we truncated all genes longer than a set truncation length *T* and calculated the average CUB of the ribosomal proteins in that genome, varying *T* from 75pb to 450pb in 75bp increments and using five different CUB metrics from the literature (Fig. 1). For the most part, CUB estimates were relatively stable up to 300bp (near the 240bp cutoff gene length *gRodon* uses by default to exclude short coding sequences [10, 25]). For sequences shorter than that, all CUB metrics showed strong biases in their estimates. Notably, these biases were directional and largely consistent across genomes, suggesting that truncation to a particular length biased estimates in a predictable way (Fig.1). Considering that short-gene CUB biases appear predictable based on gene length, we hypothesized that growth prediction models fitted on CUB estimates from short genes would yield better results when tested on short genes than would models fitted on CUB estimates from long genes.

**Figure 1:**
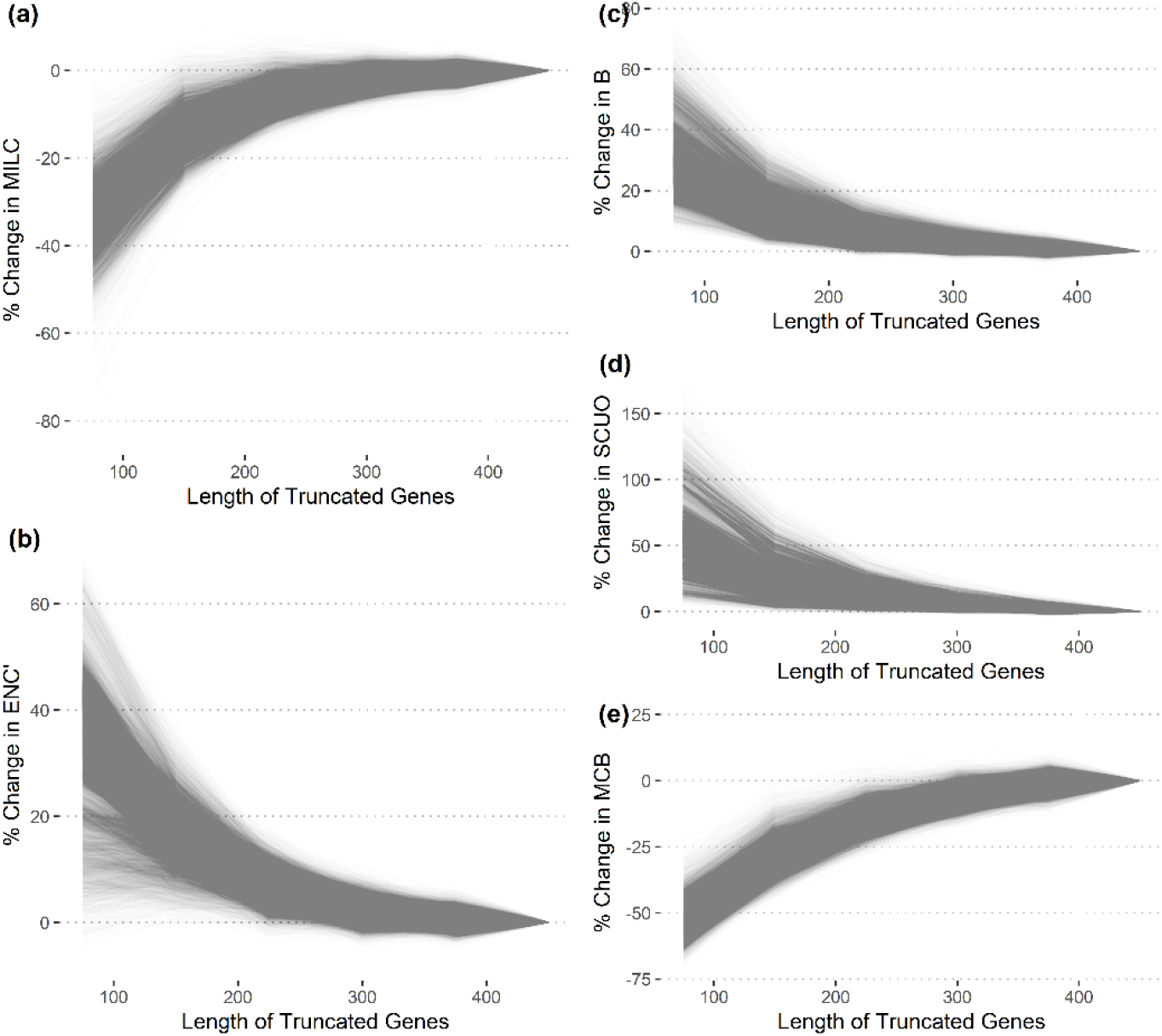
Measures of Codon Usage Bias (CUB) are unreliable for short coding sequences. Each line in each panel corresponds to a single genome’s average CUB across its ribosomal proteins where all genes in that genome were truncated to the length specified on the x-axis. Each panel corresponds to a different measure of CUB. All measures showed steep, unidirectional changes as genes were truncated to less than 240bp long (the current cutoff used for gRodon prediction). Note that in contrast to these large, monotonic changes, the overall variance of each CUB measures did not change as dramatically with truncation.

### Gene Length-Specific Models Overcome CUB Gene Length Bias

Using our progressively truncated gene sets (above), we fit linear models to predict maximum growth rates using CUB estimates from genes of various lengths (Fig. 2). Specifically, we fit linear models of log-transformed minimum doubling time using the five CUB metrics in Fig. 1 as predictors, with each model trained on CUB estimates corresponding to a different value for the truncation length *T*. Model error was minimized when the training and testing lengths shared the same *T* (Fig. 2a,b), indicating that gene length specific models outperformed models trained on full-length genes. The advantage of gene length specific models was largest for short genes, where the range of training set values of *T* that led to low model prediction error on short genes was narrower than the range of values for *T* that led to low model errors for longer genes (Fig. 2a,b). For genes longer than 300bp, any training *T* greater than or equal to 300bp led to comparable model performance. Therefore, while length specific models do not appear to be important for gene lengths longer than 300bp, they are essential for accurate maximum growth rate predictions when using genes shorter than 240bp-300bp, such as genes inferred directly from sequencing reads.

**Figure 2:**
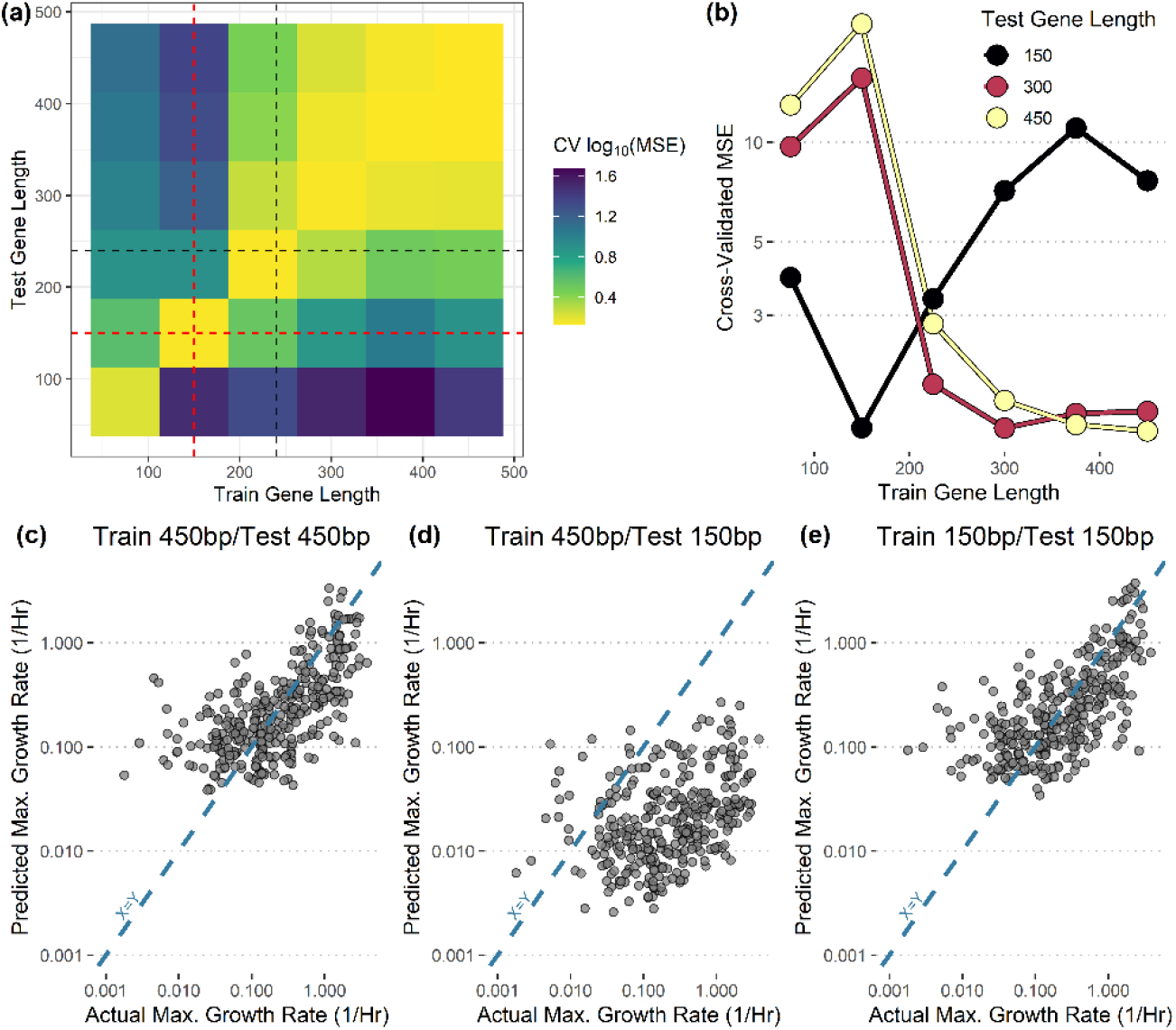
Length-specific models can accurately predict maximum growth rates from short coding sequences. (a) Model performance was highest when applied to similar gene lengths to the training data. A linear model combining the CUB measures in Figure 1 was trained and tested on CUB values taken from genomes with genes truncated to different lengths. Test-set genomes, in addition to having variable gene lengths, were iteratively held out from model training under 10-fold cross validation to yield an accurate error estimate. Red dashed lines at 150bp (a common read length) and black dashed lines at 240bp (the gRodon default cutoff) for reference. (b) Marginal plots from panel (a) for three test-lengths. Note that the error when predicting on genes truncated to 150bp is lowest when the model is trained on genes of similar length, and that this error is comparable to models trained and tested on gene lengths well above the gRodon cutoff (450bp). (c-e) Actual predictions for models trained and tested on short (150bp) and long (450bp) truncated genes.

### Applying Gene Length Specific Models to Synthetic Metagenomes

To confirm that gene length specific models also improve performance in a mixed-species metagenomic context, we assessed performance on two types of synthetic benchmarking datasets. First, we assessed model performance on “gene mixtures” truncated to different lengths *T* [17]. In this case, we simply took the complete set of genes annotated in 10 randomly selected genomes from the *gRodon* training set and combined them into one super-genome on which we performed prediction. This test captures whether gene length specific models can accurately predict the average maximum growth rate of a set of organisms when handed all of their genes at once. Second, we assessed model performance on true synthetic metagenomes [26]. In this case, from the same sets of 10 genomes we simulated realistic sequencing reads at low coverage and assuming a lognormal species abundance distribution. This test captures whether gene length specific models can find signal in the noise of a realistic dataset and cope with varying species abundances when predicting the average maximum growth rate of a community.

When applied on gene mixtures, we saw comparable results to the case of individual genomes (Fig. 3). For gene mixtures with genes truncated to 150bp to represent realistic read lengths, a model trained on 150bp genes outperformed a model trained on 450bp genes (Fig 3a,c-e). Additionally, average community predictions from the 150bp model were tightly associated with the fraction of genomes from organisms with actual doubling times <1hr in each mixture (Fig 3b; Pearson Correlation, r=-0.80, p<2.2e-16), supporting the idea that these average community-wide predictions can be thought of as a measure of the proportion of fast-growing organisms in a community.

**Figure 3:**
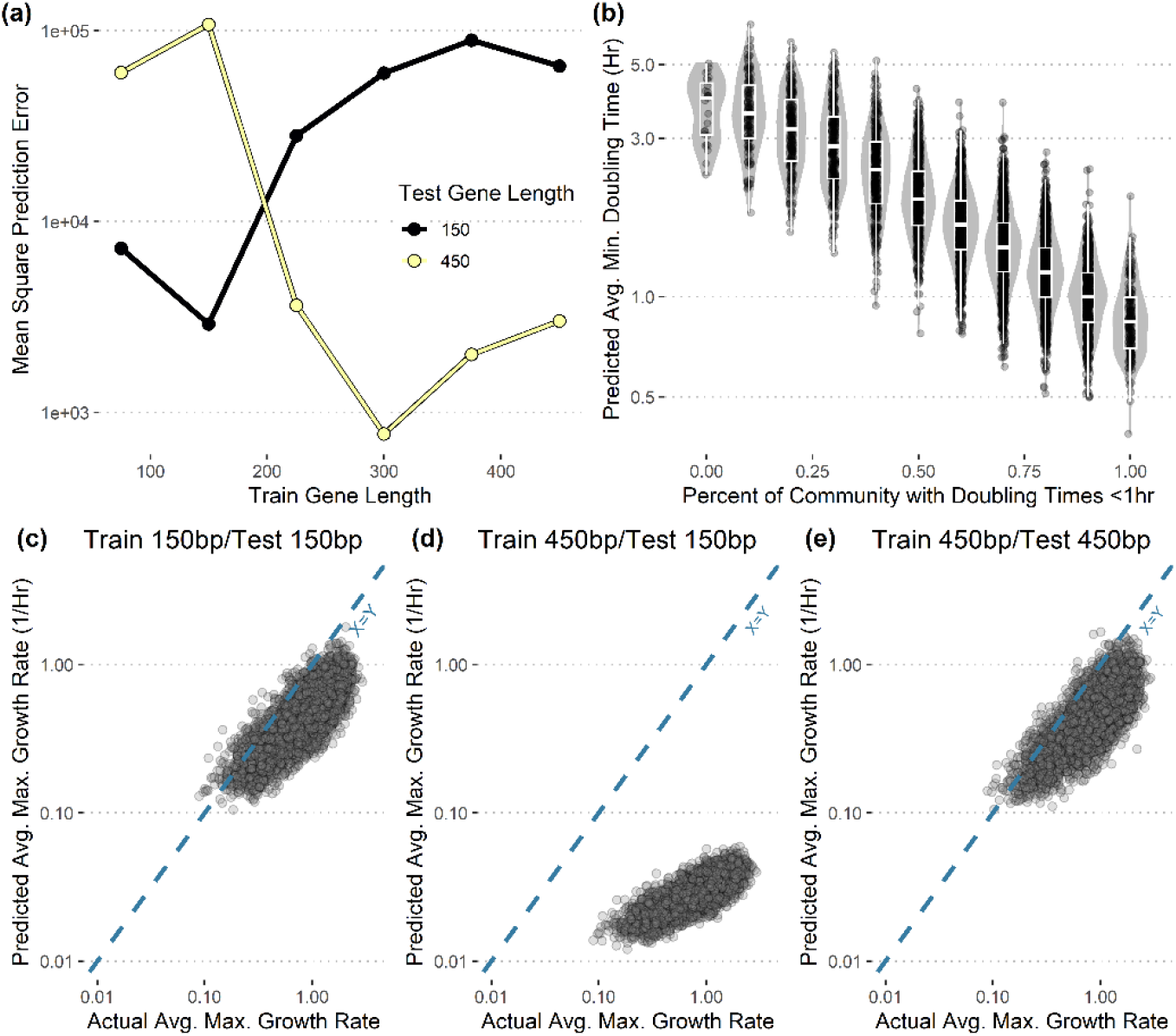
Length-specific models can accurately predict average maximum growth rates from mixtures of genomes with short coding sequences. Each genome mixture is a concatenation of 10 genomes with known maximum growth rate for each individual genome. All genes in a mixture were truncated to either 150bp or 450bp. (a) Error rates when predicting on 150bp-truncated mixtures using the 150bp model trained in Fig 2 are comparable to error rates when predicting on 450bp-truncated mixtures using the 450bp model trained in Fig 2. Alternatively, prediction on gene lengths different from model training data yields poor performance. (b) The percent of the community with doubling times <1hr (actual doubling times from the literature) is highly correlated with the average community-wide minimum doubling times predicted from genome mixtures with genes truncated to 150bp. (c-e) Actual rates and predictions for models trained and tested on short (150bp) and long (450bp) truncated genes (testing on genome mixtures; actual average calculated using a geometric mean).

Given the reliable performance of gene length specific models on mixed species assemblages, we then implemented a short read metagenomic prediction mode in *gRodon* using the model for 150bp genes. We then tested this model on genes inferred directly from simulated sequence reads from our synthetic metagenomes as well as *gRodon*’s metagenome mode v2 [17], which was trained on full-length genes. Short-read mode had lower prediction error than metagenome mode v2 (Fig 4a,c,d; Wilcoxon rank-sum test, p<2.2e-16) and predictions from short-read mode were correlated with the abundance-weighted fraction of genomes from organisms with actual doubling times <1hr in each metagenome (Pearson Correlation, r=-0.51, p<2.2e-16). Together, these benchmarks indicate that short-read mode not only outperforms existing models when applied to coding sequences inferred at the read level, but is also able to capture real patterns in growth rate variation among microbial communities even using low coverage metagenomes.

**Figure 4:**
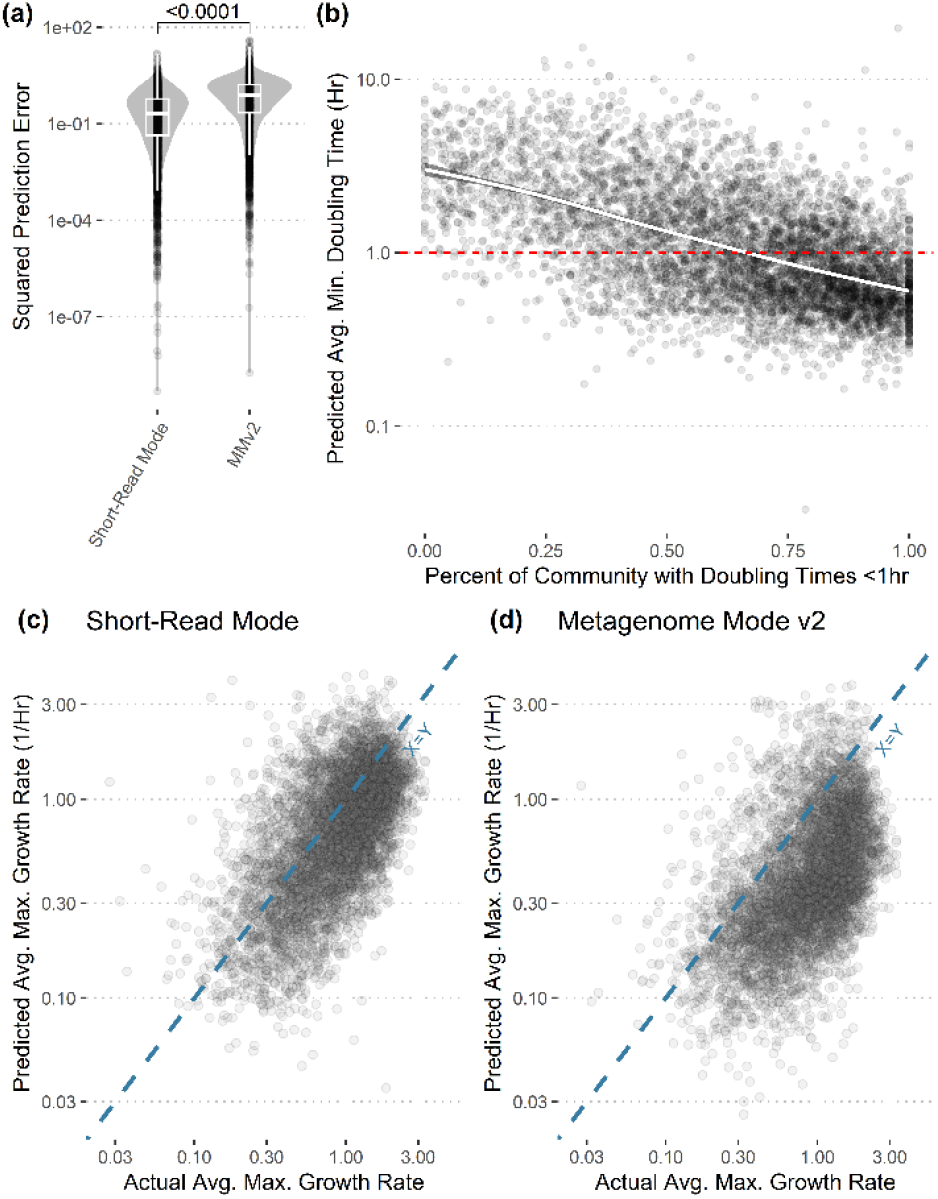
Length-specific models can accurately predict average maximum growth rates directly from reads from synthetic metagenomes. (a) Applying the short-read model trained on 150bp truncated genes, now implemented in gRodon, we found that it had lower prediction error than gRodon’s metagenome mode v2, which was trained on full-length genes (genes inferred directly from 150bp reads for synthetic metagenomes generated from the same genome mixtures as in Fig. 3 using a realistic sequencing error model and lognormal species abundances drawn at random). (b) The short-read model predictions were correlated with the overall percent of the community with fast doubling times (weighted by relative abundance). Trendline shows GAM fit. (c-d) Predictions for short-read mode and metagenome mode v2 plotted against actual average maximum growth rates (weighted geometric mean; species max. rates from literature).

## Conclusions

We developed and benchmarked a series of gene length specific maximum growth rate prediction models and showed that for short coding sequences, such as those inferred directly from sequence reads, length specific models outperform the default *gRodon* models. These new models enable assembly-free maximum growth rate prediction from metagenomes, thus allowing for community-scale metagenomic growth rate prediction even for highly complex soil communities. Even with low coverage metagenomes, length specific models are able to capture patterns in community-level maximum growth rates, meaning that expensive high coverage sequencing runs are not always needed for maximum growth rate inference [22]. We provide these models in an update to the open source *gRodon* maximum growth rate prediction tool for microbial ecologists and biogeochemical modelers hoping to integrate community growth estimation into model parameter estimation and validation (available at https://github.com/jlw-ecoevo/gRodon2; [10]).

## Methods

All scripts to generate figures can be found at https://github.com/jlw-ecoevo/gRodon-assembly-free. Short read prediction is possible using the latest version of the gRodon package available at https://github.com/jlw-ecoevo/gRodon2.

### Datasets

We performed model training and testing on the original *gRodon* training set [15] using genomes from NCBI RefSeq [26] matched to trait data gathered from the literature by Madin et al [24]. In total, this dataset represents 8,300 annotated genomes matched to trait data for 463 species of bacteria and archaea.

### Synthetic Metagenomes and Genome Mixtures

Because the set of already cultured microbial species is biased towards fast growers [15], we employed a multi-step sampling strategy to generate genome mixtures representing a range of community-level growth phenotypes. First, starting with the set of all genomes, we sampled 100 genome mixtures with 10 genomes each. Then, limited to the set of all genomes with doubling times >0.1hr, we again sampled 100 mixtures, and continued to iterate in such a way until we generated 5700 mixtures in total (the iteration went from 0.1 to 5 in increments of 0.03 in log_10_ space). This process ensured that a sizeable proportion of mixtures had slow average maximum growth rates similar to those of many natural communities.

To generate synthetic metagenomes, we first applied InSilicoSeq v2.0.1 to each mixture to generate 100k reads using the built-in novaseq error model and simulating lognormal species coverage [26]. Reads were then filtered using fastp v0.23.4 (default parameters) and coding sequences were inferred using FragGeneScanRs v1.1.0 with the illumina_1 error model [23, 28]. We annotated coding sequences as ribosomal proteins using blastn (E value cutoff of 1e−5 and a 99% identity [29]) and the original ribosomal protein database from the growthpred program for growth rate prediction [13].

### Model Training and Assessment

We trained linear regression models of log-transformed doubling times using the five CUB metrics in Fig. 1 (MILC [25], ENC’ [30], B [31], SCUO [32], and MCB [33]) as predictors. For each length specific model, CUB estimates were generated from genes truncated to the given length of that model. Model performance was assessed through a modified 10-fold cross validation procedure, where species were split into folds, CUB estimates for species in the training folds were generated from genes truncated to the training length, a model was fit, CUB estimates for species in the testing fold were generated from genes truncated to the testing length (potentially different from the training length), and the model was applied to these estimates to make a maximum growth rate prediction. This ensured we did not train and test a model on the same set of species but using different truncation lengths. The final short-read model implemented in the *gRodon* R package (Fig. 4) was fit using a Box-Cox transformation rather than a log transformation to maintain consistency with other models in the package (using the MASS R package [34]).

